# Hydrogel injection molding to generate complex cell encapsulation geometries

**DOI:** 10.1101/2021.10.31.466681

**Authors:** Amy E. Emerson, Alec B. McCall, Sarah R. Brady, Emily M. Slaby, Jessica D. Weaver

## Abstract

Biofabrication methods capable of generating complex, three-dimensional, cell-laden hydrogel geometries are often challenging technologies to implement in the clinic and scaled manufacturing processes. Hydrogel injection molding capitalizes on the reproducibility, efficiency, and scalability of the injection molding process, and we adapt this technique to biofabrication using a library of natural and synthetic hydrogels with varied crosslinking chemistries and kinetics. We use computational modeling to evaluate hydrogel library fluid dynamics within the injection molds in order to predict molding feasibility and cytocompatibility. We evaluate the reproducibility of hydrogel construct molding and extraction and establish criteria for the selection of hydrogels suitable for injection molding. We demonstrate that hydrogel injection molding is capable of generating complex three-dimensional cell-laden construct geometries using diverse hydrogel materials and that this platform is compatible with primary human islet encapsulation. We envision the application of this technique with primary cells via an off-the-shelf clinical kit that can be implemented at the patient bedside by a clinician or integrated with scaled, high-throughput automated cell manufacturing processes to generate encapsulated cell-based therapeutic products.

**Graphical Abstract:** 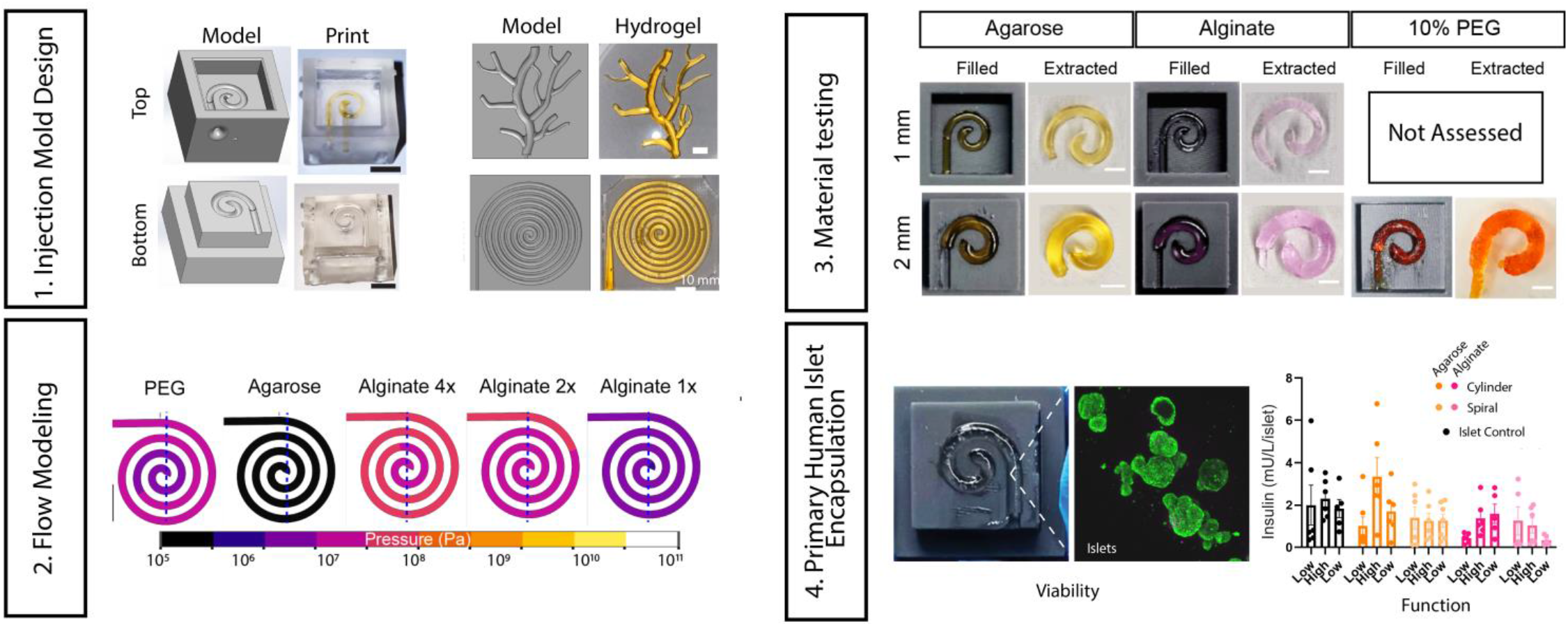

## 1. Introduction

Rapid advancements in the fields of tissue engineering and biomaterials have resulted in diverse techniques to generate complex, three-dimensional (3D) cell-containing hydrogel constructs, including methods such as 3D bioprinting and photolithography ^1^. An array of hydrogel-based materials has been adapted to these fabrication techniques, including synthetic, natural, and combination materials with a wide range of functionalities (e.g. degradable, thermoresponsive, etc.) ^2,3^. Many tissue engineering applications require hydrogel construct fabrication in anatomically-relevant shapes to regenerate specific tissues ^4,5^, whereas applications delivering cells as therapeutics (e.g. insulin-secreting cells) often require high surface-area-to-volume ratio geometries for optimal nutrient transport and therapeutic agent delivery ^6,7^.

3D bioprinting is a popular method to reproducibly fabricate complex constructs consisting of hydrogels and cells, enabling the fabrication of geometries with increasing fidelity, precision, and resolution; however, printing can take significant time and building hydrogels can be challenging ^2,3^. Additionally, 3D bioprinting faces barriers to the clinic due to complex equipment, setup, and operation requirements, and the likely need for specialized technicians or training; this particularly limits the accessibility of this technique in low-resource settings ^8^. 3D bioprinting is also challenging to scale efficiently in manufacturing, with added complexity to maintain sterility and implement clinical good manufacturing practices ^9^. Therefore, there is a pressing need for an easier, faster, and more reliable way to encapsulate cells in complex hydrogel geometries, both accessibly in the clinic and efficiently in encapsulated cell product manufacturing.

Injection molding is a widely used process to efficiently generate reproducible products in scaled and automated manufacturing using a wide array of materials ^10^. In plastic injection molding, for example, plastic precursor solutions are injected with high pressure into molds to form complex geometries with high fidelity and high reproducibility ^11^. To date, complex molding techniques have shown promise with hydrogels alone ^12^, but injection molding has not been exploited for the generation of complex cell-laden hydrogel geometries in the clinic or in scaled cell product manufacturing processes. In applications where a single hydrogel and cell type are required in a complex geometry, injection molding has the potential to be an efficient, intuitive, and reproducible approach to generate complex hydrogel constructs. While the typical intra-mold pressures associated with injection molding are incompatible with biofabrication and cell survival, computational modeling-aided mold design can optimize device characteristics to produce comparable flow regimes and localized mechanical forces, and therefore comparable cytocompatibility, to biofabrication techniques like extrusion bioprinting ^9,13,14^. Additionally, studies have shown that hydrogels can shield cells from shear stresses during flow, an additional protective factor during injection molding biofabrication ^15^.

Biofabrication techniques with the broadest applications are those that are compatible with a versatile array of hydrogels ^2,3,16,17^. Injection molding has comparable hydrogel characteristic requirements to 3D bioprinting: (1) hydrogels must support cell viability during construct fabrication, (2) pre-gelation solutions must be capable of injection or flow, and (3) post-gelation hydrogel mechanical strength must be sufficient to maintain the construct geometry ^3,17^. Common cell-supportive hydrogels used in biofabrication techniques include natural hydrogels, such as alginate and agarose, and synthetic hydrogels, such as poly(ethylene glycol) (PEG) ^2^, although there are a wide range of natural, synthetic, and combination materials adapted for use in biofabrication techniques^16,17^.

We report an injection molding-based method to generate cell-encapsulating hydrogels with complex geometries at the patient bedside (**Figure 1A**). We demonstrate the versatile geometries and range of hydrogels compatible with injection molding, including the high surface area-to-volume spiral geometry (**Figure 1B**) designed for the specific application of islet macroencapsulation as a cell therapy to treat type 1 diabetes. Macroencapsulation of primary islets enhances their retrievability *in vivo*, while maintaining a high surface area-to-volume ratio geometry maximizes nutrient transport for optimal cell survival and function ^7^. We model fluid dynamics within spiral injection molds and characterize the relationship between reproducibility of hydrogel fabrication and extraction and hydrogel mechanical properties. Last, we characterize the impact of injection molding on the viability and function of a model beta cell line and primary human islets. We ultimately envision the dissemination of this technology via commercially available clinical hydrogel injection molding kits for a wide range of hydrogel and/or cell therapy applications (**Figure 1C**), as well as for scaled, high-throughput automated manufacturing of encapsulated cell products. Additionally, this technology is accessible to any laboratory with access to a 3D printer and computer-aided design (CAD) software.

**Figure 1.**
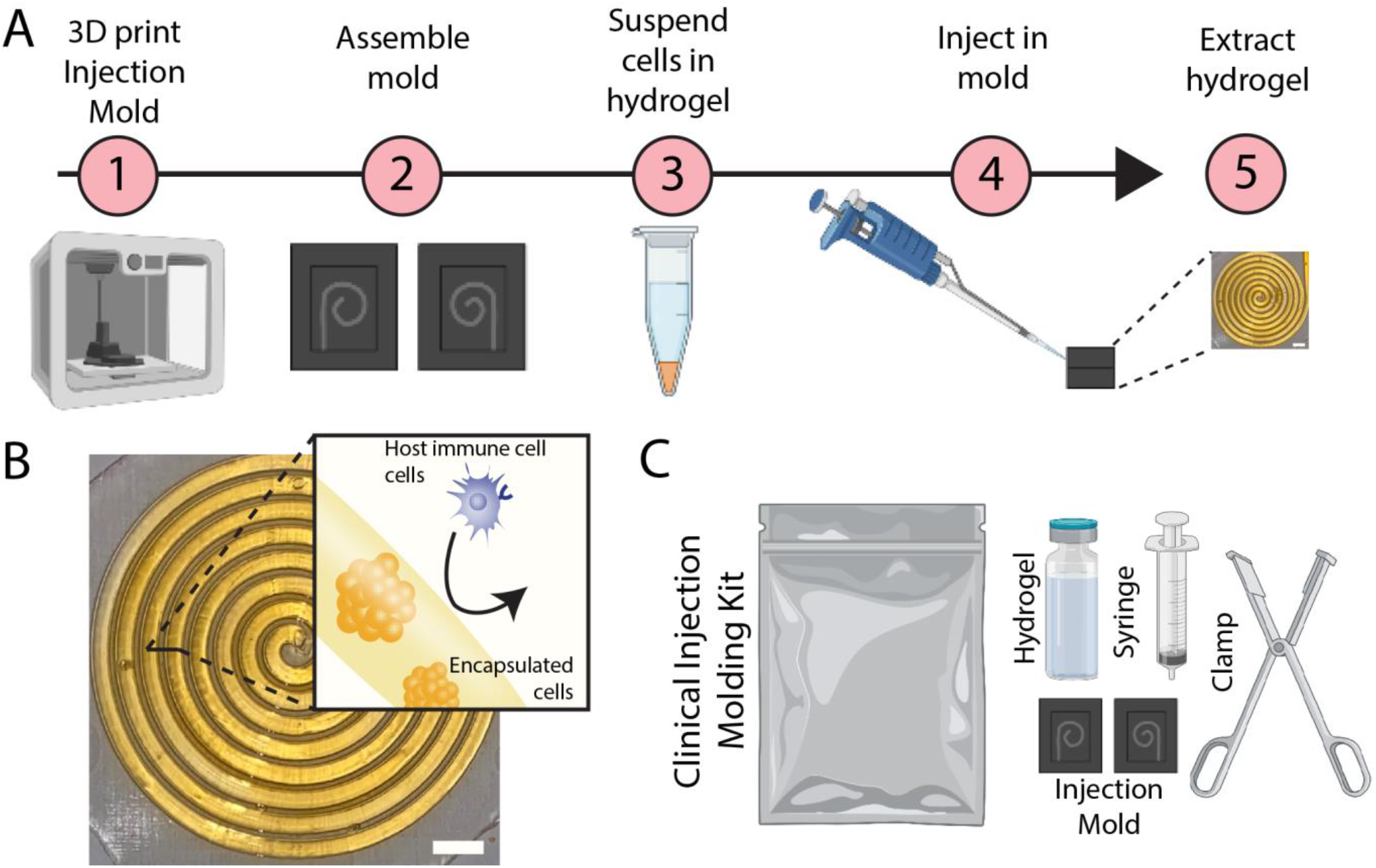
Workflow and components of a hydrogel injection molding-based cell macroencapsulation method. (**A**) To generate hydrogels of complex geometries, injection molds are 3D printed, assembled, and injected with a mixture of hydrogel and cells. Complex geometries, such as a spiral (**B**), can be generated in degradable or nondegradable hydrogels to deliver encapsulated cells, such as islets. (**C**) Hydrogel and injection molding components can be delivered in a sterile kit for use by surgeons in the clinic. Scale bar = 5 mm. Figure designed with the help of Biorender.com

## 2. Materials and methods

### 2.1 Materials

Chemicals were obtained from Sigma Aldrich (St. Louis, MO) and cell culture materials were obtained from Thermo Fisher (Carlsbad, CA) unless otherwise noted. Peptides were obtained from Genscript (Piscataway, NJ) unless otherwise noted. The Flexible and Clear Resins compatible with the Formlabs Form 3 3D printer were obtained from Formlabs (Somerville, MA).

### 2.2 Solidworks injection mold design and flow simulation

Injection mold devices were designed in Solidworks with a set tolerance dependent on the selected resin. The tolerance used for the primary resin (flexible) was 0.05 mm. Solidworks Computational Fluid Dynamics Flow Simulation, which solves the Navier-Stokes equations, was conducted to evaluate static pressure and velocity of hydrogels injected into the mold. Volumetric flow rate into the mold was set at 30 µL/s. Flow simulation internal analyses were conducted for alginate (2% w/v), agarose (2% w/v), and PEG (5 and 10% w/v) with viscosity (μ) of each gel measured in its pre-gelation state. Density (ρ) values were obtained from the literature. For alginate (2% w/v), μ = 24.79 Pa*s and ρ = 1010 kg/m^3 18^. For agarose (2% w/v), μ = 1.129 Pa*s and ρ = 1010 kg/m^3^. For PEG μ = 75 Pa*s and ρ = 1016 kg/m^3 19^. Viscosity of pre-gelled solutions was measured via rheometry, taking the average values at 1 minute post-initiation of gelation conditions.

### 2.3 3D printing of injection molds and mold assembly

Injection mold devices were 3D printed on the Formlabs Form 3 stereolithography 3D printer. A flexible resin was used for printing the devices used experimentally. The highest resolution, 50 µm, was used for printing due to the small dimension of the designs within the mold. Finished prints were washed for 10 minutes in an isopropyl alcohol (70%) bath, then air dried and finally cured in a Formlabs post-curing oven for 15 minutes at 60 °C. To compress injection mold halves together and form a seal, we designed and printed a custom adjustable clamp 3D printed with Formlabs Tough 2000 resin.

### 2.4 PEG-maleimide hydrogels

20 kDa four-arm poly(ethylene glycol) (PEG)-maleimide powder (Laysan Bio, Arab, AL) was combined with sterile Dulbecco’s phosphate buffer solution without magnesium chloride and calcium chloride (DPBS^-/-^) in a 0.25 mg/µL solution and a 0.125 mg/µL for a 10% PEG solution and 5% PEG solution, respectively (pH 7.0-7.4). Dithiothreitol (DTT) powder was combined with DPBS^-/-^ at a concentration 0.035 mg/µL and 0.00175 mg/µL for each PEG solution, respectively. Both solutions were vortexed until homogenous. The PEG and DTT solutions were combined in a 1:1 ratio to form PEG hydrogels, with polymer and crosslinking solutions injected sequentially into the molds.

### 2.5 Agarose hydrogels

Low-gelling temperature agarose powder was combined with sterile DPBS with magnesium chloride and calcium chloride (DPBS^+/+^) (2% w/v) in a 50 mL beaker in a double boiler. The solution was stirred with a stir bar at 600 rpm and 100 °C until completely homogeneous. For visualization of hydrogels in photography, a small volume of orange food dye (U.S. Art Supply, San Diego, CA) was added.

### 2.6 Slow-gelling alginate hydrogels

Slow-gelling alginate was fabricated using alginate (UP-MVG, Novomatrix), calcium carbonate (CaCO_3_) and gluconic acid δ-lactone (GDL), where varied concentrations of GDL and CaCO_3_ enable the tuning of crosslinking rate (**Table 1**). Alginate and GDL solutions were made in DPBS^-/-^ and stored at 4 °C for 24 hours to allow for full solubilization. After 24 hours, CaCO_3_ was added to the alginate solution and mixed well, and finally the GDL solution was added in a 1:1 ratio to form the final alginate (2% w/v) mixture. For photography purposes, 3 drops of pink food dye (U.S. Art Supply, San Diego, CA) were added to the GDL solution before gelation.

**Table 1.** Composition of slow-gelling alginates of various gelation speeds.

|  | CaCO<sub>3</sub>:GDL<br>(mM:mM) |
| --- | --- |
| Alginate 1x | 15:30 |
| Alginate 2x | 30:60 |
| Alginate 4x | 60:120 |

### 2.7 Characterization of hydrogel properties via rheometry

Hydrogel storage and loss moduli describing the linear viscoelastic region of hydrogels were assessed by strain and frequency sweeps performed at 25 °C on a Physica MCR 101 rheometer (Anton Paar, Austria) with a 25 mm sandblasted parallel plate geometry. The hydrogels for mechanical characterization (490 µL, n=3/group) were synthesized and swollen overnight in DPBS^+/+^ (agarose) or DPBS^-/-^ (alginate) before placement. The system was lowered to a gap distance ranging from 1.3 mm (agarose), 1.4-1.6 mm (alginate), 1.0-1.4 mm (PEG) depending on the minimum distance necessary for adequate normal force detection, as gel height will vary depending on material used. Strain amplitude sweeps (0.01% to 1000% strain) were conducted at 1 Hz to characterize the linear viscoelastic region. A constant strain value of 0.1% strain for frequency sweeps (0.1 Hz to 100 Hz) was chosen based on this region.

Time studies were conducted to assess the gelation point via storage and loss modulus crossover for hydrogels. The hydrogel solutions were mixed and immediately injected onto the rheometer at 25 °C. Hydrogel storage and loss modulus were recorded over time (0.1% strain, 1 Hz) to display the crossover gelation point, as well as the complex viscosity of the sample before and after gelation; agarose measurements started at 100 °C and was allowed to cool to 25 °C.

### 2.8 Injection mold reproducibility and extraction studies

For 1 mm and 2 mm diameter spirals, 30 µL and 75 µL of gel was injected into clamped 3D printed molds, respectively. Excess hydrogel was used to ensure complete filling. The hydrogel solution was set in the mold for up to 10 minutes. Reproducibility of fill was measured by injecting the set volume of gel into the injection molds and measuring the percent fill. After injection and a short incubation period to allow gelation, the clamp and top half of the injection mold were removed, and the hydrogel extracted. Images of hydrogels in and out of the mold were taken for analysis using a camera mounted on a custom frame. A custom FIJI program cropped, scaled, and thresholded the gel inside of the injection mold to measure the cross-sectional area of gel in mm^2^. The measured area was normalized to a 100% filled spiral. Reproducibility of extraction was measured as the percent of successfully fabricated whole gels extracted completely from the mold (n = 10/group).

### 2.9 Cell encapsulation and assessment in injection molded hydrogels

Alginate (2% w/v) gels and agarose (2% w/v) gels were fabricated as described above. The beta cell line INS-1E, kindly gifted by Andres J. Garcia, was cultured in RPMI supplemented with 10% FBS, 1% penicillin-streptomycin, 1% HEPES buffer solution, 1% Sodium Pyruvate, and 0.09% beta-mercaptoethanol. INS-1E cells were passaged, counted, and aliquoted to ensure a uniform cellular density of 7500 cells/µL (to correspond to a density of 5 islet equivalents/µL) for each hydrogel cylinder and spiral. The cells were suspended in hydrogel solutions, injected into the injection mold or open cylinder mold, and allowed to gel for 5 minutes. The hydrogel constructs were carefully removed from their molds and cultured in well plates under standard conditions (5% CO_2_, 20% O_2_). At 24 hours post-construct fabrication, the alamar blue assay was mixed in a 1:10 ratio with the complete RPMI media per the manufacturer instructions and cultured with hydrogels for 4 hours prior to fluorescence measurement (exc/em 560/590 nm). The resulting data was normalized to hydrogel volume and the control hydrogel. Live/dead (Calcein AM/ethidium homodimer, 30 minutes) confocal imaging was performed at 0 and 24 hours. Images were collected with a Leica SP8 White Light Laser Confocal microscope, housed in the Regenerative Medicine and Bioimaging Facility at Arizona State University, and analyzed using ImageJ. Percent live/dead was calculated by dividing the area of live/green fluorescence by the total area of fluorescence. All images were thresholded with the same values.

### 2.10. Islet encapsulation and assessment in injection molded hydrogels

Alginate (2% w/v) and agarose (2% w/v) gels were fabricated as described above in 1 mm diameter spiral injection molds, cylindrical control hydrogels, or cultured in standard 2D conditions. Human islets (Prodo Labs) were received and cultured overnight in standard conditions (5% CO_2_, 20% O_2_) in pancreatic islet culture medium (Prodo Labs, Aliso Viejo, CA) at a density of 50-100 IEQ/mm^2^ surface area. The islets were encapsulated 24 hours post-receiving at a density of 20 IEQ/µL and cultured 48 hours prior to assessment. Glucose stimulated insulin response (GSIR) was performed in static incubation and standard culture conditions in KREBS buffer (15 mM NaCl, 4.7 mM KCl, 1.2 mM MgSO_4_, 2.5 mM CaCl_2_, 26 mM NaHCO_3_, and 0.2% BSA) with low (3 mM) and high (11 mM) glucose concentrations. Encapsulated or unmodified islets were exposed to low glucose buffer for a 1-hour pre-incubation, and two hour sequential low-high-low incubations. Samples were frozen at -80° C for later analysis via human insulin ELISA (Mercodia, Uppsala, Sweden). Alamar Blue metabolic assay was performed according to manufacturer instructions and cultured with samples for 4 hours prior to fluorescent measurement. The data was normalized to hydrogel volume (i.e. cell number) and control hydrogel values. Live/dead (calcein AM/ethidium homodimer) staining was performed per manufacturer instructions and imaged via confocal microscopy. Confocal image live/dead percentages were analyzed using ImageJ; all images were thresholded with the same values.

### 2.11 Statistics

Hydrogel precursor solution viscosities were analyzed by one-way ANOVA with Kruskal-Wallis multiple comparisons. Mold filling was analyzed by unpaired t-test and one-way ANOVA with Holm-Sidak’s multiple comparison test for 1 and 2 mm spirals, respectively. Storage modulus and time to critical viscosity was analyzed by one-way ANOVA with Kruskal-Wallis multiple comparison test against 5% PEG or agarose, respectively, (Figure 4) or against every group (Supplementary Figure 2). In single cell experiments, the percent live cells were analyzed using an ordinary two-way ANOVA with Sidak’s multiple comparison between alginate and agarose materials. Alamar blue data was analyzed using a two-way ANOVA. In human islet experiments, GSIR data was analyzed with one-way ANOVA. Live cell percentage, Alamar blue, and index values were analyzed by non-parametric one-way ANOVA with Kruskal-Wallis multiple comparisons against control islets.

## 3. Results

### 3.1. Design and fabrication of complex hydrogel geometries via injection molding

We used a 3D stereolithography system to print injection molds capable of generating hydrogel constructs with complex geometry (**Figure 1**). 3D printed methacrylate resin-based injection molds were assembled with a custom clamp to ensure a tight seal, and a hydrogel, with or without cells, was injected into the injection mold using a pipette or syringe (**Figure 1A**). Hydrogel injection molding can potentially be performed with a wide range of hydrogels, requiring only that the material crosslinking time is slow enough to enable mixing and injection and sufficient hydrogel mechanical stiffness to withstand removal from the mold. The injection molding method is intuitive and requires minimal training, and we ultimately envision an off-the-shelf kit that can be implemented bedside during surgery, consisting of an injection mold, a hemostat-inspired customized clamp, and hydrogel components (**Figure 1C**). Optional primary cells (e.g. pancreatic islets) can be incorporated by the clinician at the point of delivery.

Injection molds were designed using computer-aided design (CAD) software and 3D printed using a flexible methacrylate-based printing material (**Figure 2A**). Stereolithography printing enables the generation of injection molds in a variety of methacrylate-based materials, including opaque materials of a variety of stiffnesses and clear materials for visualization of the injection molding process (**Figure 2A**). We chose the “flexible” resin for reproducibility and cell encapsulation experiments due to its slight flexibility, superiority in generating a tight seal for injection molding, and capacity to withstand autoclave sterilization. Additionally, this resin has a print resolution of 50 µm to 0.5 mm. To ensure a tight seal and prevent hydrogel leakage prior to crosslinking, we designed a customized adjustable clamp (**Figure 2B**) modeled after an adjustable wrench for intuitive use. Preliminary testing with agarose hydrogels demonstrated that injection molding can generate a versatile selection of complex geometries (**Figure 2C-D**) including spirals with a resolution down to 0.5 mm arm width (**Figure 2C)** and geometries with more complex flow patterns and sharp corners (**Figure 2D**). Spirals were chosen as the primary injection molded geometry for future experiments due to their favorable surface area-to-volume ratio, which should provide efficient transport of oxygen and nutrients to encapsulated cells ^7^.

**Figure 2.**
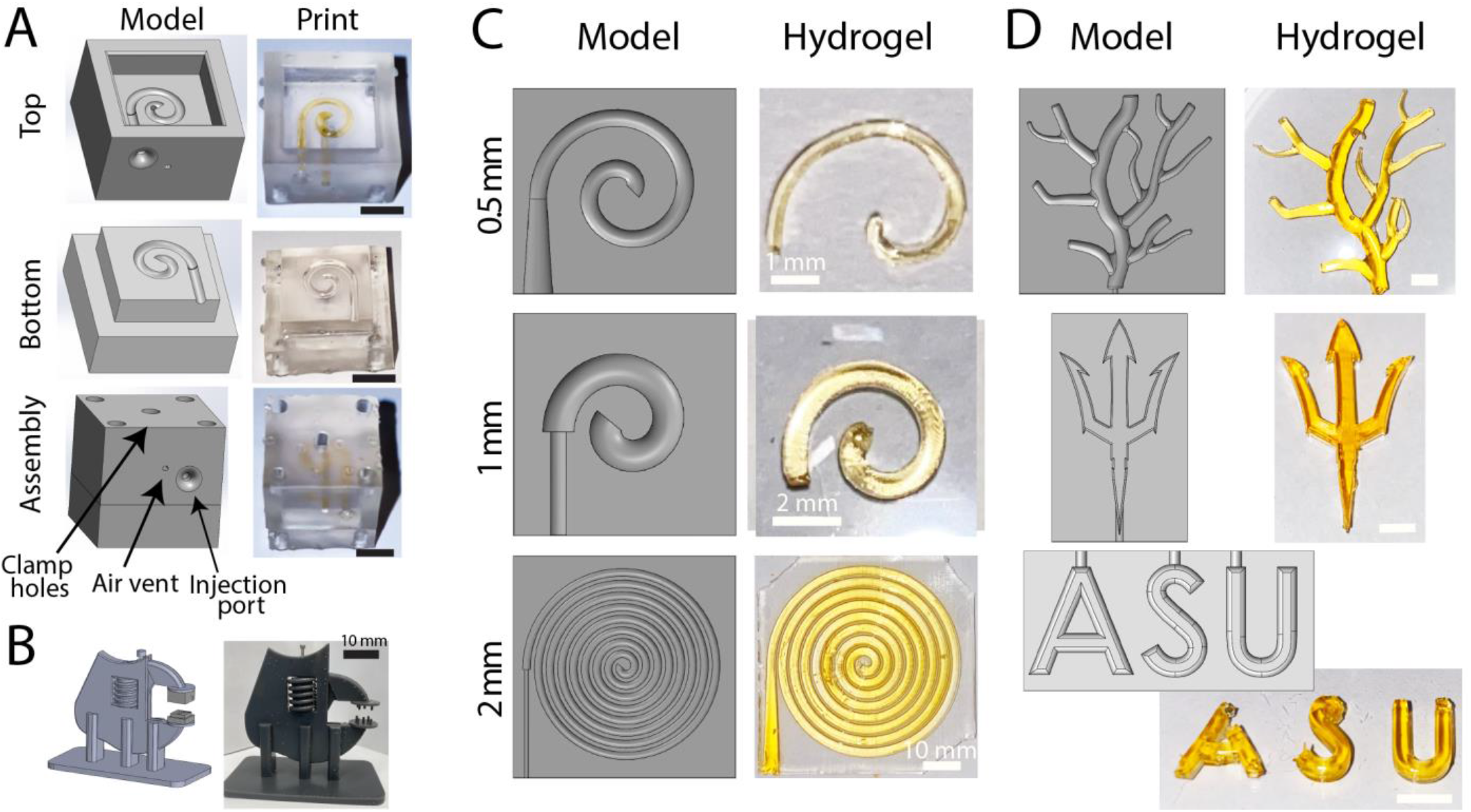
Generation of complex hydrogel geometries via an injection molding system. (**A**) CAD models of injection mold parts are printed on a stereolithographic 3D printer, shown here in a clear resin to visualize print details, including injected agarose (orange). The mold is held together via an adjustable clamp during crosslinking (**B)**. To demonstrate the versatility of agarose geometries generated using injection molding, we tested various spiral lengths and widths (0.5 – 2 mm spiral arm diameter) (**C**), as well as more complex shapes (**D**), such as blood vessels and the Arizona State University trident and logo. All scale bars 5 mm unless otherwise noted.

### 3.2. Modeling hydrogel flow within injection mold devices to predict molding feasibility and cytocompatibility

We next sought to evaluate the influence of hydrogel viscosity and injection mold geometric shape on pressure, velocity, and flow profiles within spiral injection molds via computational fluid dynamics simulations (**Figure 3**). Given the potential for cytotoxicity at pressure levels above 200 MPa ^20^, we were interested in identifying the maximum pressure experienced in spiral injection molds designed for preclinical (i.e. rodent) and clinical (i.e. human) scale hydrogel encapsulation device designs. Computational flow simulations can predict forces experienced by cells in the injection molds *in silico*, as well as predict the feasibility of generating a complex geometry. We developed a library of hydrogel materials (**Table 2**) to evaluate in injection molding with a range of crosslinking kinetics, pre-gelled solution viscosities, and mechanical properties.

**Table 2.** Properties of hydrogels evaluated in flow modeling and injection molding.

| Sample | Hydrogel Stiffness (Pa) | Complex viscosity<sup>+</sup> (Pa*s) |
| --- | --- | --- |
| PEG 5% | 50 | NA |
| PEG 10% | 301 | 75 |
| Alginate 1x | 1293 | 25 |
| Alginate 2x | 4867 | 183 |
| Alginate 4x | 4967 | 333 |
| Agarose | 5483 (25°C) | 1.75 (100°C) |

NA = Not Assessed, <sup>+</sup> measured at one min post initiation of gelation

**Figure 3.**
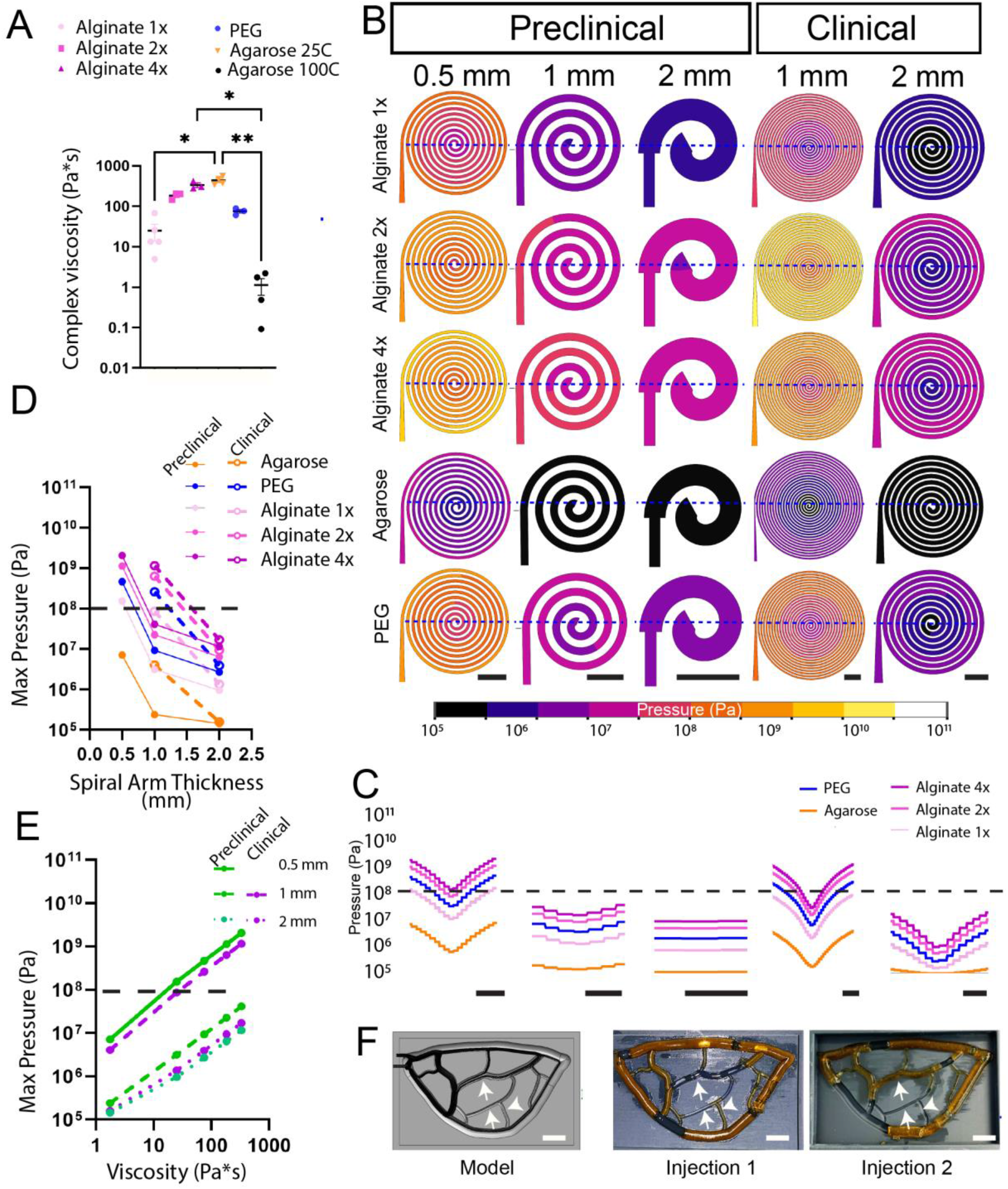
Modeling hydrogel fluid flow dynamics in injection molds. Complex viscosity measurements (**A**) of PEG, alginate, and agarose solutions pre-gelation enabled modeling of hydrogel pressure (Pa) dynamics within injection molds spatially (**B**) and quantitatively with respect to geometry (**C**) and location within the geometry (**D**) using a volume flow rate of 30 µL/s. Plotting viscosity against injection molding pressure (**E**) illustrates the influence of viscosity on pressure. (**F**) Flow modeling within the devices (black lines) is predictive of injection molding success for agarose in a capillary bed injection mold. Dashed lines in **C, D**, and **E** indicate threshold of cytotoxic pressure (∼10^8^ Pa). White arrows indicate areas the model predicted would result in low pressure and reduced flow. Scale bars = 5 mm. Error = SEM. Viscosities analyzed by one-way ANOVA with Kruskal-Wallis multiple comparisons. * P < 0.05, ^**^ P < 0.01

We performed time-dependent rheological measurements on pre-gelled 2% alginate (1-4x, see methods and **Table 1**), 2% agarose (at 25°C and 100°C), and 10% PEG (w/v) solutions to obtain viscosity values at one minute post initiation of gelation (**Figure 3A**). One-minute post-initiation of gelation was presumed to be the maximum time a typical user would need to inject the mixture into the device, and we assumed an estimated volume flow rate of 30 µL/s. Given agarose’s temperature-dependence and rapid gelation speed, resulting in significantly different viscosities at different temperatures (**Figure 3A**), we used the dynamic viscosity measurement for warm agarose (**Table 2**) under the presumption that the agarose solution is kept warm until just prior to injection. Static pressure (**Figure 3B-D)** and velocity (**Supplementary Figure 1**) were modeled in the Solidworks computational fluid dynamics module for alginate, agarose, and PEG hydrogels within spiral geometries of varied arm diameter (0.5 to 2 mm) and scale (preclinical or clinical sizes). The greatest pressures were experienced at the spiral inlet and were highest in longer spirals with smaller diameter arms, peaking at greater than 10^9^ Pa for 0.5 mm diameter arms (**Figure 3C** and **D**). The alginate solutions, which possess a higher viscosity than pre-gelation agarose, displayed 10-fold higher pressure values than agarose across all geometries. Plotting maximum pressure against viscosity illustrates the linear relationship between these variables and their dependence on geometry (**Figure 3E**). Velocity remained consistent between materials, and only varied with construct geometry (**Supplementary Figure 1**). Finally, flow simulations generally predicted the feasibility of fabricating more complex geometries within the injection molds. In one example of flow modeling predicting failure, a capillary bed-type geometric design displayed flow trajectories that indicate weak flow areas (white arrows) that were then observed as defects in two experimental trials (**Figure 3F)**.

### 3.3. Impact of hydrogel mechanical properties on injection molding feasibility

A critical feature of a clinical hydrogel injection molding device is the reproducibility and completeness of hydrogel construct molding and extraction. The variables expected to impact these outcomes the greatest are hydrogel gelation times and hydrogel mechanical properties. As such, we selected a library of hydrogel materials to test in our injection molding platform: varied weight percentages of synthetic (PEG)-based hydrogels as well as naturally derived matrices alginate and agarose. This material selection provides a range of material properties and crosslinking times to investigate the constraints of our hydrogel injection molding method.

First, rheometry was used to measure time-dependent hydrogel complex viscosity (**Figure 4A**), with agarose and alginate 4x demonstrating the fastest approach to the viscosity cutoff (gray region, > 300 Pa*s for the 1 mm preclinical spiral injection mold, as modeled in **Figure 3B-E**). 10% (w/v) PEG hydrogels, which crosslink via the rapid maleimide-thiol Michael addition reaction, and alginate 1x hydrogels each reached the cutoff viscosity at 13.3 min, comparable kinetics to alginate 2x (**Figure 4B**).

**Figure 4.**
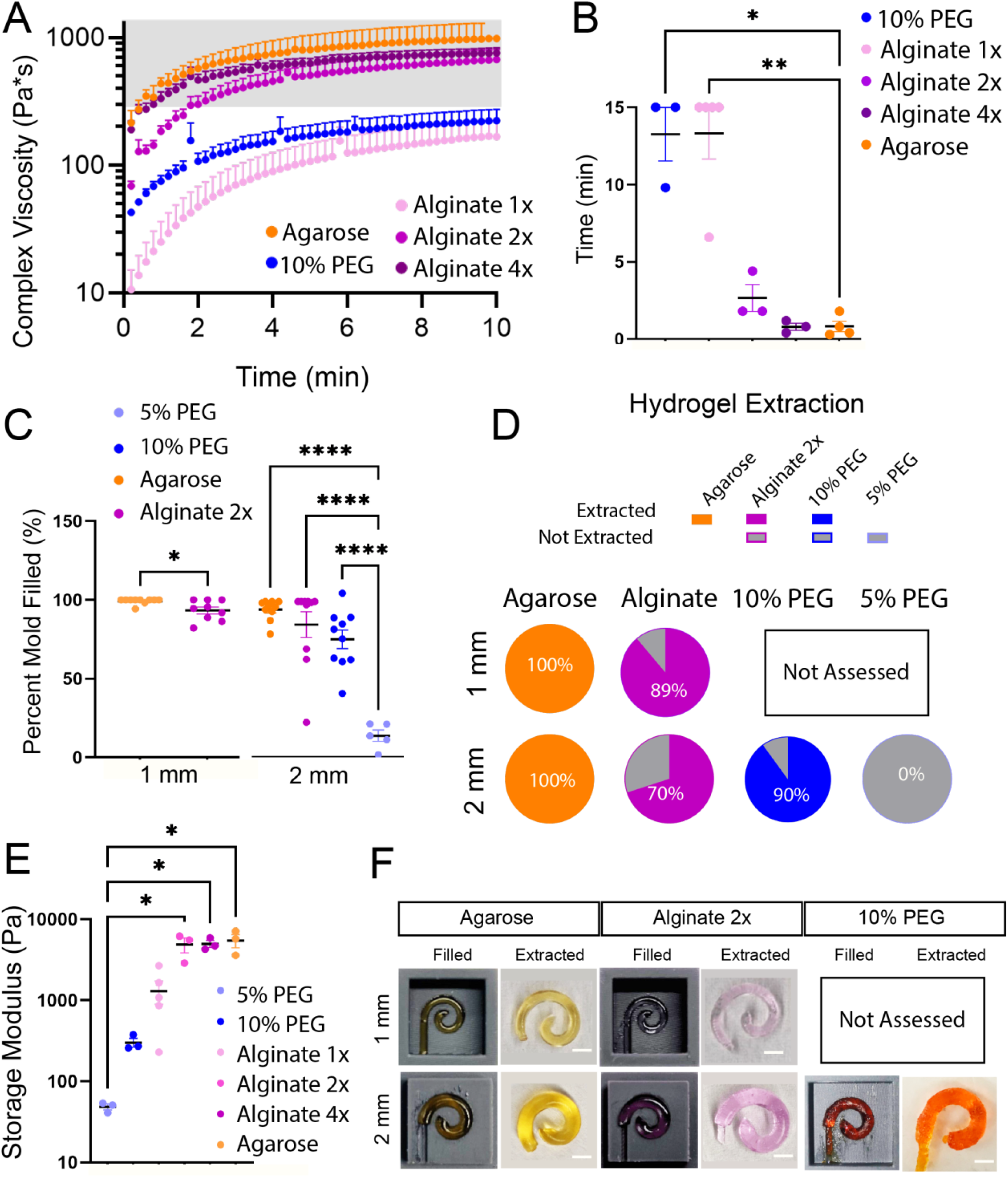
Hydrogel properties influence suitability for injection molding. (**A**) Alginate, agarose, and PEG hydrogels vary in time to reach the critical viscosity of 300 Pa*s (gray region), as quantified in (**B**). We tested the reproducibility of injection mold complete filling (**C**) for spirals injected with PEG, alginate and agarose for 1 and 2 mm diameter spirals. The percent of gels able to be completely extracted (**D**) varied greatly by material, likely governed by material mechanical stiffness post-gelation (**E**). (**F**) Example images of filled molds and successfully extracted PEG, alginate, and agarose hydrogels. Scale bars = 2 mm. Error = SEM. Mold filling analyzed by unpaired t-test and one-way ANOVA with Holm-Sidak’s multiple comparison test for 1 and 2 mm spirals, respectively. Storage modulus and time to critical viscosity analyzed by one-way ANOVA with Kruskal-Wallis multiple comparison test against 5% PEG or agarose, respectively. * P < 0.05, ^**^ P < 0.01, ^****^ P < 0.0001

We next investigated the reproducibility of injection mold hydrogel filling and extraction using PEG, agarose, and alginate hydrogels. Alginate 2x was chosen due to its moderate crosslinking time relative to other alginate formulations and moderately high stiffness (4867 Pa) post-crosslinking; alginate 1-4x exhibited a trend (p = 0.0054) of increasing mechanical stiffness (1300, 4867, and 4967 Pa for 1x, 2x, and 4x alginate, respectively), though individual groups were not significant from each other (**Supplementary Figure 2**). The completeness of mold filling is critically dependent on hydrogel crosslinking time, as hydrogels that crosslink too quickly cannot flow easily through the mold. We first measured the percent area of filled cavity within the mold using image analysis. We quantified the filled area (**Figure 4C**), finding that agarose and alginate averaged above 93% filled area for 1 mm diameter spirals and above 85% filled area for 2 mm spirals. We also attempted to inject PEG hydrogels (5 and 10% w/v) in 2 mm spiral molds and found reduced completeness of mold filling with both PEG gel concentrations relative to alginate and agarose, though only 5% PEG exhibited significant reductions in completeness (14% filled area). There was no significant difference between the agarose and alginate percent fill for the 2 mm diameter spiral.

Last, we investigated the reproducibility of hydrogel full extraction from the injection molds, measured based on the percent of moldings that resulted in the extraction of complete spiral hydrogels (**Figure 4D**). All agarose hydrogels (100%) were extracted from 1 and 2 mm diameter spirals (n = 9-10), whereas 89%, 70%, and 90% of 1 mm alginate spirals, 2 mm alginate spirals, and 10% PEG hydrogels were successfully extracted, respectively (n = 10). PEG hydrogels with a lower polymer concentration were challenging to extract from the molds, with 0% of the 5% PEG gels extracted, (n = 9). We evaluated the mechanical stiffness of materials in our hydrogel library via rheometry (**Figure 4E**) and found that stiffer hydrogels, such as agarose (storage modulus 5483 Pa), were most readily and completely extracted from the device, whereas decreasing mechanical stiffness reduced the probability of success (alginate 2x and 10% PEG with storage moduli of 4867 and 301 Pa, respectively). The material with the lowest stiffness, 5% PEG (50 Pa), exhibited the poorest probability of extraction, at 0%. Representative images showing mold filling and spiral extraction are shown in **Figure 4F**.

### 3.4. Influence of injection molding on cell viability

As our modeling of pressure levels within the spiral injection molds indicated a significant, and potentially cytotoxic level ^20^ of pressure during injection molding, particularly in thinner spiral arms and more viscous solutions (**Figure 3B**), it is critical to evaluate the influence of pressures within the mold on encapsulated cells and validate our theoretical cytotoxic pressure cutoff point. We evaluated 1 and 2 mm spiral arm diameters against a traditionally-molded hydrogel cylinder control construct (see **Supplementary Table 1** for dimensions) for both agarose and alginate (2% w/v) hydrogels. Open-molded cylindrical hydrogels are expected to experience only standard ambient pressure.

We encapsulated the model beta cell line INS-1E to evaluate cell survival via live/dead imaging (**Figure 5A**) and percent live cell quantification (**Figure 5B-D**) at 0 and 24 hours post-molding, as well as metabolic activity via alamar blue assay at 24 hours (**Figure 5E**). Cells were encapsulated at a density of 7500 cells/µL (the equivalent to 5 primary islet equivalents (IEQ) per µL) in all encapsulation devices. Immediately post-injection, cells in agarose constructs exhibited greater than 90% viability regardless of injection mold geometry, whereas more viscous alginate solutions exhibited a decline in cell viability to 87% and 76% for 2 and 1 mm geometries, respectively (**Figure 5B**). By 24 hr post-injection molding, alginate and agarose constructs maintained comparable viability to the 0 hr timepoint, apart from 1 mm agarose spirals, which exhibited reduced viability (84%). Across both hydrogels, 1 mm spiral geometries exhibited a significant (p < 0.0001) reduction in cell viability relative to cylindrical controls. Additionally, alginate demonstrated consistently reduced viability relative to agarose injection molded hydrogels across both time points (**Figure 5C**). By plotting the percent viability of each construct at 24 hr post-injection (**Figure 5B)** against the maximum pressure we predicted cells would experience during fabrication of each geometry (**Figure 3B-D**), we can observe the relationship between ambient pressure within our devices and cell survival (**Figure 5D**). Assessment of cell metabolic activity at 24 hr post-encapsulation via alamar blue demonstrated that the 1 mm spiral hydrogels had 2.2 and 1.7 fold greater metabolic activity than the cylinders for agarose and alginate constructs, respectively (**Figure 5E**). For both alginate and agarose, 2 mm spiral constructs had comparable metabolic activity to cylinders.

**Figure 5.**
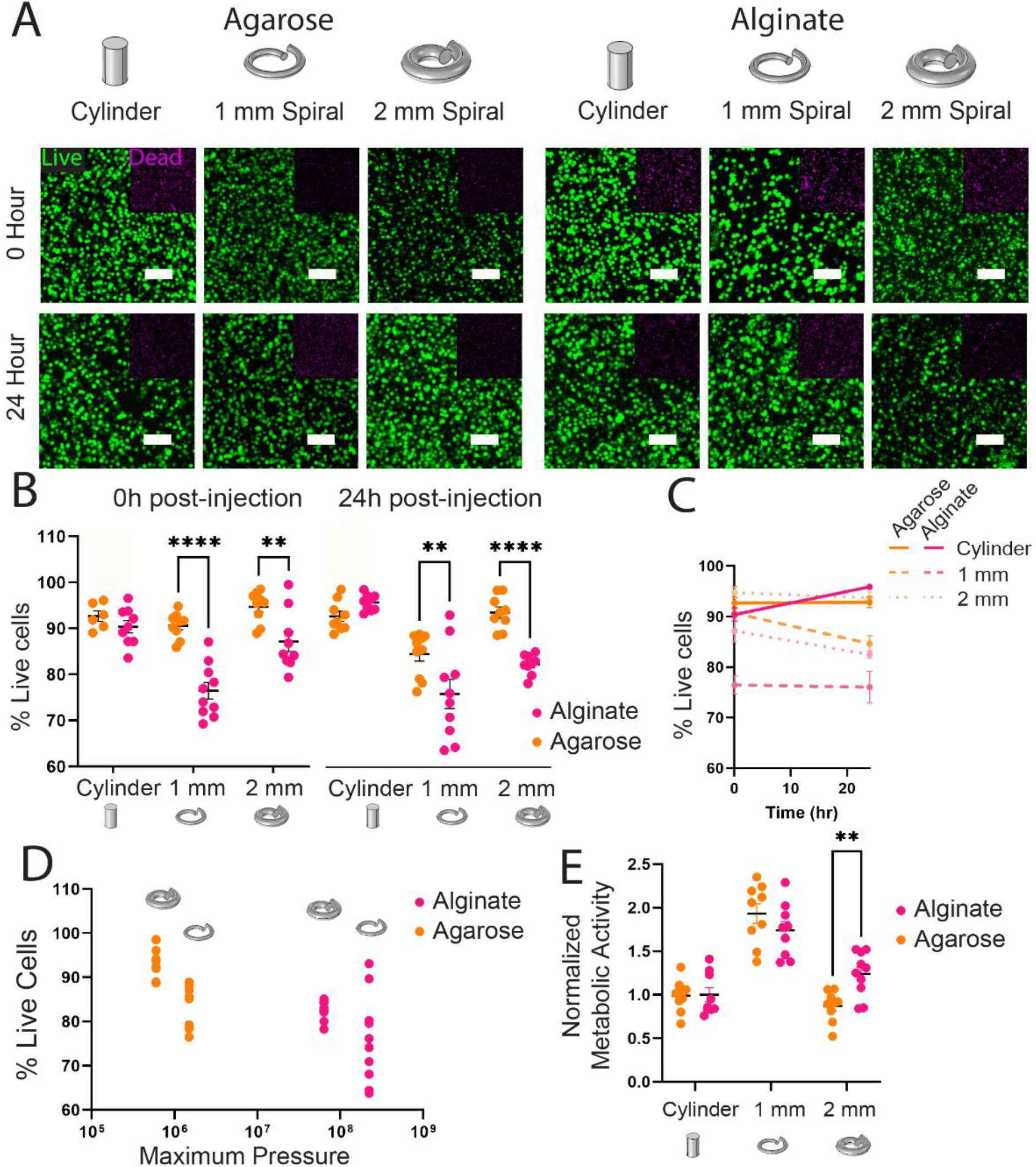
Influence of injection molding on encapsulated model beta cell line viability and metabolic activity. (**A**) Live (green)/dead (magenta) confocal imaging of the INS-1E beta cell line at 0 and 24 hours post-molding in cylinder or 1 or 2 mm spiral geometries formed in either agarose or alginate hydrogels. (**B**) Live cell % was quantified (black line = mean) from the images in (**A**) (n = 6-10/group) and plotted against time in (**C**). 24 hours viability data plotted against the average pressure experienced within the injection mold for that material and geometry in (**D**). Cell metabolic activity was evaluated at 24 hours post-molding (n = 9/group) (black line = mean) (**E**). Each data point represents an individual gel, data was pooled from three independent experiments. Live/dead and alamar blue data was analyzed via one-way ANOVA with Sidak’s multiple comparison. Error bars = SEM. Scale bars = 100 µm. ^****^ P < 0.0001, ^**^ P < 0.01, * P < 0.05.

### 3.5 Influence of injection molding process on primary human islet viability and function

Given the potential detrimental effects observed in a robust and proliferative model cell line post-injection molding (**Figure 5B**) and the intention for this device to be implemented with primary cells in the clinic, it was critical to evaluate the impact of injection molding on the viability and function of non-proliferative primary human islet cells (**Figure 6**). As in our model cell line experiments, we evaluated the influence of spiral injection molding (1 mm arm diameter) on encapsulated human islet viability and function against an open molded hydrogel cylinder control geometry in both alginate and agarose hydrogels (2% w/v), and an unencapsulated islet control group. Islets were encapsulated at a density of 20 IEQ/µL.

**Figure 6.**
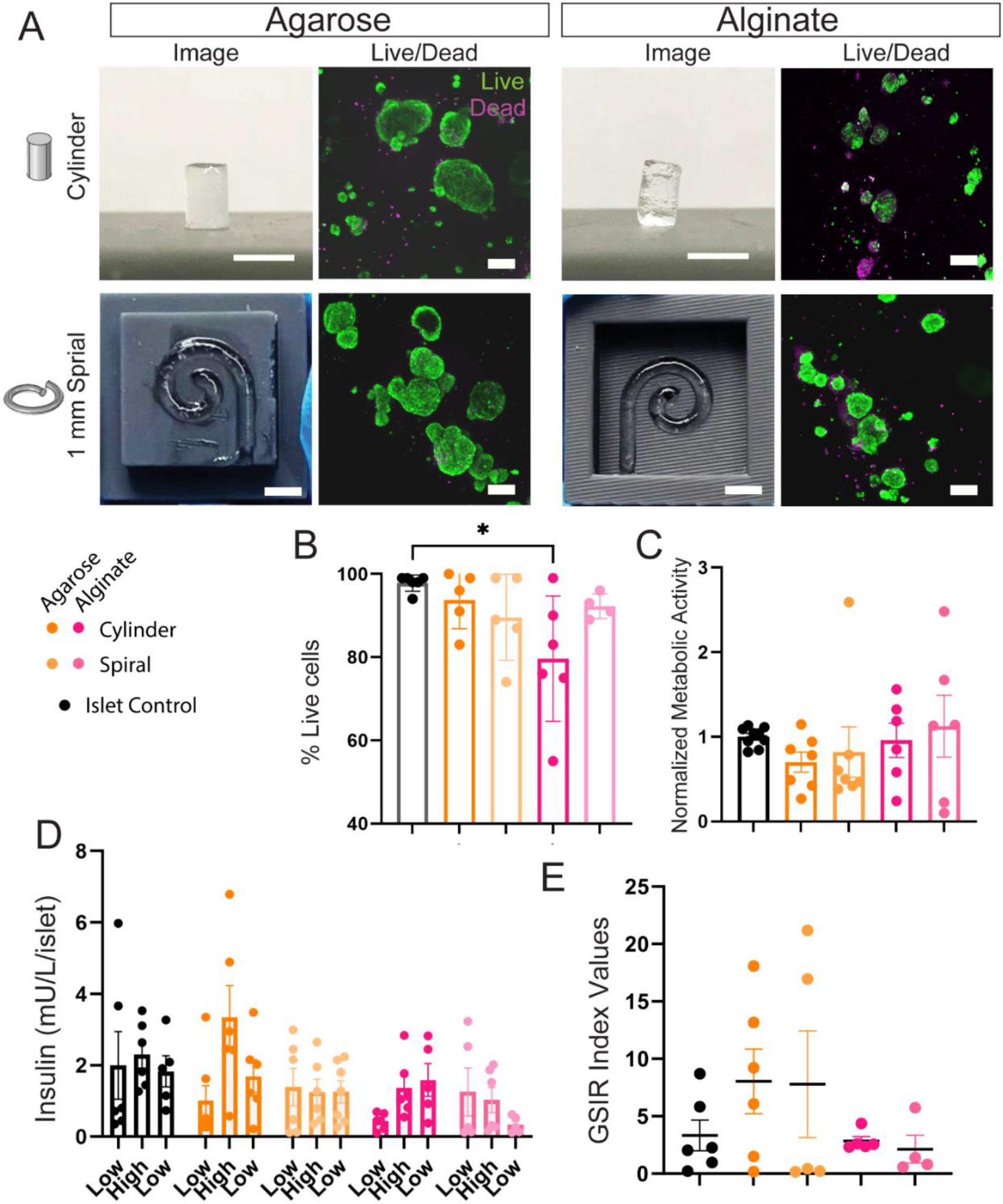
Human islet viability and function after injection molding. (**A)** Live (green)/dead (magenta) confocal imaging of human islets encapsulated in 1 mm spiral or cylindrical control geometries 48 hr post-molding, in either agarose or alginate hydrogels. (**B**) Percent viable cells quantified from live/dead imaging at 48 hr. At 48 hr post molding, metabolic activity (**C**) of the encapsulated cells was evaluated via alamar blue assay, and insulin responsiveness was assessed by glucose-stimulated insulin response (GSIR) assay (**D**). (**E**) GSIR index values, the ratio of high glucose to low glucose insulin secretion. n = 4-6 gels/group A-B, n = 6-7 gels per group C-E. Replicates were pooled from three independent experiments. GSIR data analyzed by one-way ANOVA. % Live cells, alamar blue, and index values analyzed by non-parametric one-way ANOVA with Kruskal-Wallis multiple comparisons against control islets. * P < 0.05. Error bars = SEM. Scale bars: live/dead = 200 µm, macroscopic images = 5 mm.

Live/dead confocal imaging of human islets within the macroencapsulation devices (**Figure 6A**) demonstrated a high degree of islet viability across both geometry and hydrogel configurations (**Figure 6B**), with only the alginate cylinder group exhibiting significantly reduced islet viability (∼20% reduction). Islet metabolic activity (evaluated via alamar blue) demonstrated comparable values across all groups at 48 hours post-molding (**Figure 6C**; individual human islet batch data in **Supplementary Figure 3**). Additionally, we evaluated human islet insulin responsiveness via the Glucose Stimulated Insulin Response (GSIR) assay, where low (3 mM), high (11 mM), and second low (3 mM) glucose incubations should elicit a characteristic biphasic insulin response (**Figure 6D**). Three separate preparations of human islets from non-diabetic donors were pooled, resulting in a high degree of variation in GSIR data. An additional measure of insulin responsiveness is the GSIR index value (high glucose value normalized to first low glucose value); all groups exhibited moderate average index values of 2.1-8.0, with no significant differences in insulin responsiveness (**Figure 6E**).

## 4. Discussion

With the rise of cell-based therapies as next-generation therapeutics, there is a pressing need to develop translational cell encapsulation and hydrogel-based cell delivery methods ^21^. Cell products delivering a therapeutic, such as insulin for type 1 diabetes ^22^ or FVIII to treat hemophilia A ^23^, would particularly benefit from delivery in an immune-isolating hydrogel to (1) reduce the need for chronic systemic immunosuppression and its host of detrimental side effects ^7,24^; and (2) enhance cell product retrievability for optimum patient safety ^25,26^. Traditional cell encapsulation techniques generate small microgels ^24^, which are challenging to reliably retrieve, particularly from the traditional intraperitoneal cavity transplantation site ^27^. Macroencapsulation devices which house larger cell numbers can enhance graft safety and retrievability, but often suffer from detrimental oxygen gradients due to their scale ^7,27^. Therefore, cell macroencapsulation devices require geometric designs that prioritize oxygen transport profiles ^27^, while remaining feasible to fabricate in the clinic.

The injection molding cell encapsulation method is intuitive and requires minimal training, and we ultimately envision an off-the-shelf kit that can be implemented bedside during surgery, consisting of an injection mold, a hemostat-inspired customized clamp, and hydrogel components (**Figure 1C**); primary cells (e.g. pancreatic islets) can be incorporated by the clinician at the point of delivery. We anticipate that the simplicity of this technique could enable encapsulated cell transplantation even in low-resource settings.

In addition to accessible clinical cell delivery methods, with the rise of methods to scale and manufacture cell-based products ^28–31^, there is also a need to develop cell encapsulation methods amenable to scaled, high-throughput encapsulated cell product manufacturing ^28^. Depending on application, these cell delivery constructs often require complex three-dimensional structures (**Figure 2**) that are challenging to fabricate in a high-throughput manner via biofabrication methods, such as 3D bioprinting ^9^. Injection molding as a technique is particularly amenable to scaled manufacturing, as evidenced by its ubiquitous use in high throughput and automated manufacturing of plastic products ^10^. There are some limitations to the complexity of the 3D geometry achievable through hydrogel injection molding, although there are potential workarounds through the incorporation of sacrificial materials and via creative injection mold engineering.

We modeled static pressure (**Figure 3**) to simulate the forces that cells could experience as they travel through the injection mold. The physics of fluid flow within the molds is governed by the Navier-Stokes equations, where the relevant variables are solution viscosity, velocity, and mold geometry ^32^. Gel precursor solution viscosity (**Figure 3A**) can be used to modulate static pressure within the injection molds (**Figure 3B-D**). Larger molecular weight alginates, for example, have higher viscosities than lower molecular weight alginates at the same solution concentration ^33^. Lower viscosity solutions reduce localized pressure within the device during injection molding (**Figure 3E**) and enable faster injection speeds or smaller mold features without increasing pressure. While 2% alginate solutions were used in this study for the sake of simplicity, typical concentrations in islet encapsulation range from 1-3% ^34^; therefore, lower alginate concentration solutions have reduced viscosities, and thus lower static pressure. Additional ways to mitigate pressure include reducing hydrogel precursor velocities via programmable syringe pumps in an automated, high-throughput encapsulated cell manufacturing workflow, and modulating the air vent hole size to reduce back pressure at the mold outlet.

A particular advantage of the injection molding method is its compatibility and reproducibility with diverse hydrogels, as long as they meet two primary criteria: (1) maintains sub-critical complex viscosity long enough to allow hydrogel injection, and (2) has sufficient mechanical stiffness (>100 Pa) post-gelation to maintain hydrogel shape and facilitate removal from the mold (**Figure 4**). We investigated materials widely used in cell encapsulation and biofabrication: natural materials agarose and alginate ^2,24^, and synthetic poly(ethylene glycol) (PEG)-based hydrogels ^7,35^. This material library uses a range of injection molding-compatible crosslinking strategies: thermal, ionic, and covalent for agarose, alginate, and PEG hydrogels, respectively.

A limitation of the agarose and PEG-maleimide hydrogels is their rapid crosslinking kinetics, requiring that they be injected fairly quickly (estimated in models at 30 µL/s) ^36^. The PEG-maleimide (4-arm, 20 kDa) system uses Michael addition chemistry to create covalent bonds with dithiolated crosslinkers ^7,24,37^. This reaction occurs rapidly at physiological pH but the speed can be reduced by increasing pH ^36^. While time dependent rheometry indicated that PEG hydrogels reach the critical viscosity for 1 mm spiral devices by ∼13 minutes post initiation of gelation (**Figure 4A**), the heterogeneity of crosslinking results in a non-injectable solution within seconds. Similarly, thermoresponsive agarose gels quickly (< 1 minute, **Figure 4A**) at room temperature once removed from heat. The resultant high injection speeds generate relatively high pressures within the device during injection, ranging from a peak of 10^9^ and low of 10^5^ Pa for 0.5 mm and 2 mm diameter spirals, respectively (**Figure 3B-D**). Therefore, extending hydrogel crosslinking times by modulating hydrogel crosslinking chemistry should improve reproducibility and reduce user error.

Alginate is typically crosslinked by divalent ions that diffuse in from an external crosslinking solution. To generate a slow-gelling alginate, calcium was liberated from calcium carbonate via disassociation by gluconic acid δ-lactone (GDL) in varied concentrations (alginate 1-4x, **Table** 1, see methods for details). While alginate, agarose, and 10% PEG hydrogels readily met the crosslinking kinetics and mechanical stiffness requirements for injection molding, 5% PEG hydrogels with lower mechanical stiffness (50 Pa) were challenging to extract from the molds (**Figure 4D**). Finally, the hydrogel geometry also influences feasibility of extraction; larger spirals become unwieldy to manipulate and remove from the molds without losing their shape. Complex geometries will need to account for hydrogel manipulation and transfer in geometry design.

Previous studies have demonstrated irreversible detrimental effects on eukaryotic cell viability and function at static pressures broadly in the range of 100-200 MPa ^20^. While the injection molding devices used for cell encapsulation in this study (**Figure 5** & **6**) did not exhibit modeled pressures in this range (**Figure 3**), greater cell death (20-30% cell death) was observed in spiral devices relative to non-injection molded encapsulated cells (< 10% cell death) both immediately after and 24 hours after encapsulation, indicating the molding process itself may cause cell damage. Interestingly, the higher viscosity alginate demonstrated significantly more cell death than agarose (**Figure 5A-C**), which aligns with our models indicating higher pressure observed for this more viscous material (**Figure 5D**). We do not expect significant cell death contributions from shear stress, as previous work has demonstrated that cell injection in hydrogel carriers are likely protected relative to cells injected in solution, particularly in shear-thinning materials such as alginate ^15^. Finally, we evaluated encapsulated cell metabolic activity to confirm viability observations and were surprised to find that the constructs exhibiting the greatest cell death (1 mm spirals) exhibited higher metabolic activity 24 hours post-encapsulation (**Figure 5D**). There are two potential reasons why this assay would conflict with viability data: (1) cells are in a heightened state of stress, resulting in higher metabolic activity in injection molded groups, or (2) the higher surface area:volume ratio in the 1 mm spiral enables more rapid diffusion of the metabolic indicator into the surrounding solution relative to the other geometries.

Due to the observation of a potentially cytotoxic pressure level, we had some concern that poorly proliferative primary islets, with a limited capacity for self-repair ^38^, could experience exacerbated cytotoxic effects during injection molding. Additionally, islets are cell clusters, which could experience shearing and disassociation during flow through the molds ^39^. We used primary human cadaveric islets to evaluate the clinical translatability of injection molding-based encapsulation, as human islets are less robust to environmental stress relative to rodent islet preparations and demonstrate a higher degree of variability due to greater donor diversity ^40^. Additionally, human islets are less insulin responsive than rodent primary islets (index values of 10 vs. 30, respectively ^41^) and human islets become progressively more dysfunctional over time in culture ^42^. The human islet preparations used in this study were shipped between 4 and 7 days post isolation; as insulin responsiveness declines after 5-7 days *in vitro* culture, control islets were not highly responsive (index value 3.33, **Figure 6D**). While donor variability **(Supplementary Figure 3**) limited our ability to discern whether human islet functionality is preserved post-encapsulation, it is evident that islet viability (**Figure 6A-B**) and function (**Figure 6C-E**) was not substantially impeded by the molding process, given there were no statistical differences between encapsulated and unencapsulated control islet viability and function measures. The significant decrease in viability observed in alginate cylinders in **Figure 6B** could be partly due to imaging artifacts, as the hydrogel cylinders’ dimensions make them difficult to image deeply. Additionally, no islet organoid shearing or accelerated death was evident in injection molded groups (**Figure 6A**).

While we modeled both preclinical and clinical (human) scale device pressure and velocity (**Figure 3, Supplementary Figure 1**), cell viability and function testing focused on preclinical-scale devices to facilitate preclinical rodent studies and conserve donated cadaveric human islets. Future studies will validate the impact of modeled flow and pressure on islet viability and function in clinical scale devices. Additionally, future studies will address challenges in clinical scale device fabrication; while 2 mm clinical scale spiral devices were feasible to fabricate, they are unwieldy to extract from the mold without a sacrificial hydrogel cast over the spiral for stability (**Figure 2C**). In the application of encapsulated islet transplantation, we anticipate casting a vasculogenic hydrogel within the negative spaces of the spiral to encourage enhanced vascularization, and therefore oxygen availability, at the device surface *in vivo* ^7,24,37^. This should provide sufficient stability to manipulate larger, less stable spiral geometries.

## 5. Conclusions

Herein, we demonstrate that hydrogel injection molding is suitable to generate complex three-dimensional cell-laden construct geometries using diverse hydrogel materials. We establish criteria to develop a hydrogel library compatible with injection molding at the clinical and manufacturing scale and demonstrate the feasibility of this platform in the application of primary human islet encapsulation. We envision the application of this technique with primary cells via an off-the-shelf clinical kit that can be implemented at the patient bedside by a clinician. Further, with the development of cell-based therapy products, we envision the implementation of injection molding in the cell product manufacturing process, to generate scaled, high-throughput automated manufacturing of encapsulated cell-based therapeutics.

## 6. Acknowledgements

This research was supported by the Juvenile Diabetes Research Foundation (grant 1-INO-2020-915-A-N) and the Arizona Biomedical Research Centre New Investigator Award. The confocal microscope used in these studies was acquired by an NIH SIG award (1 S10 RR027154-01A1) and is housed in the Regenerative Medicine Imaging Facility at Arizona State University. We would also like to acknowledge the Arizona State University SciHub for the use of its 3D printer, which was used for preliminary test prints.

## 8. Author contributions

J.D.W, A.B.M., and A.E.E. developed the concept and design. J.D.W, A.B.M., and A.E.E. designed the study and analyzed data. A.B.M., A.E.E., S.R.B., and E.M.S. conducted experiments and data collection, and J.D.W. assisted with data analysis. All authors contributed to writing the manuscript.

## 10. Supplementary Figures

**Supplementary Table 1.** Dimensions of constructs used in experiments in Figures 5 and 6

| Geometry | Cylinder radius (mm) | Height/length (mm) | Volume (μL) |
| --- | --- | --- | --- |
| 1 mm spiral | 0.5 | 19.7 | 15.5 |
| 2 mm spiral | 1.0 | 18.0 | 56.7 |
| cylinder | 1.5 | 5.0 | 35.3 |

**Supplementary Figure 1.**
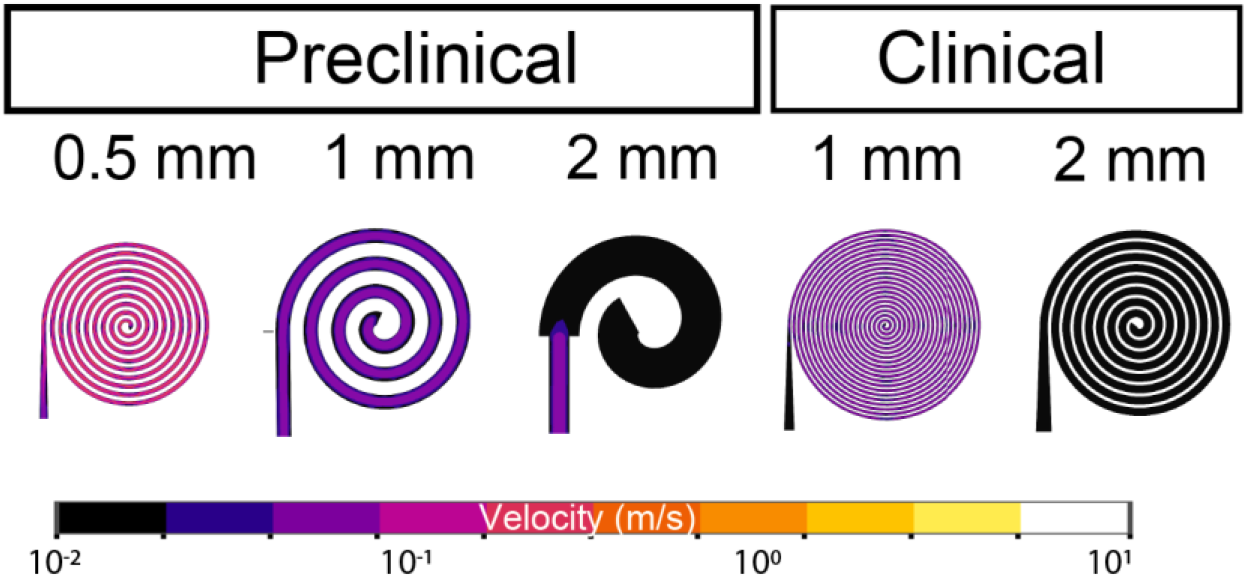
Velocity profiles of different hydrogel shapes within the injection mold.

**Supplementary Figure 2.**
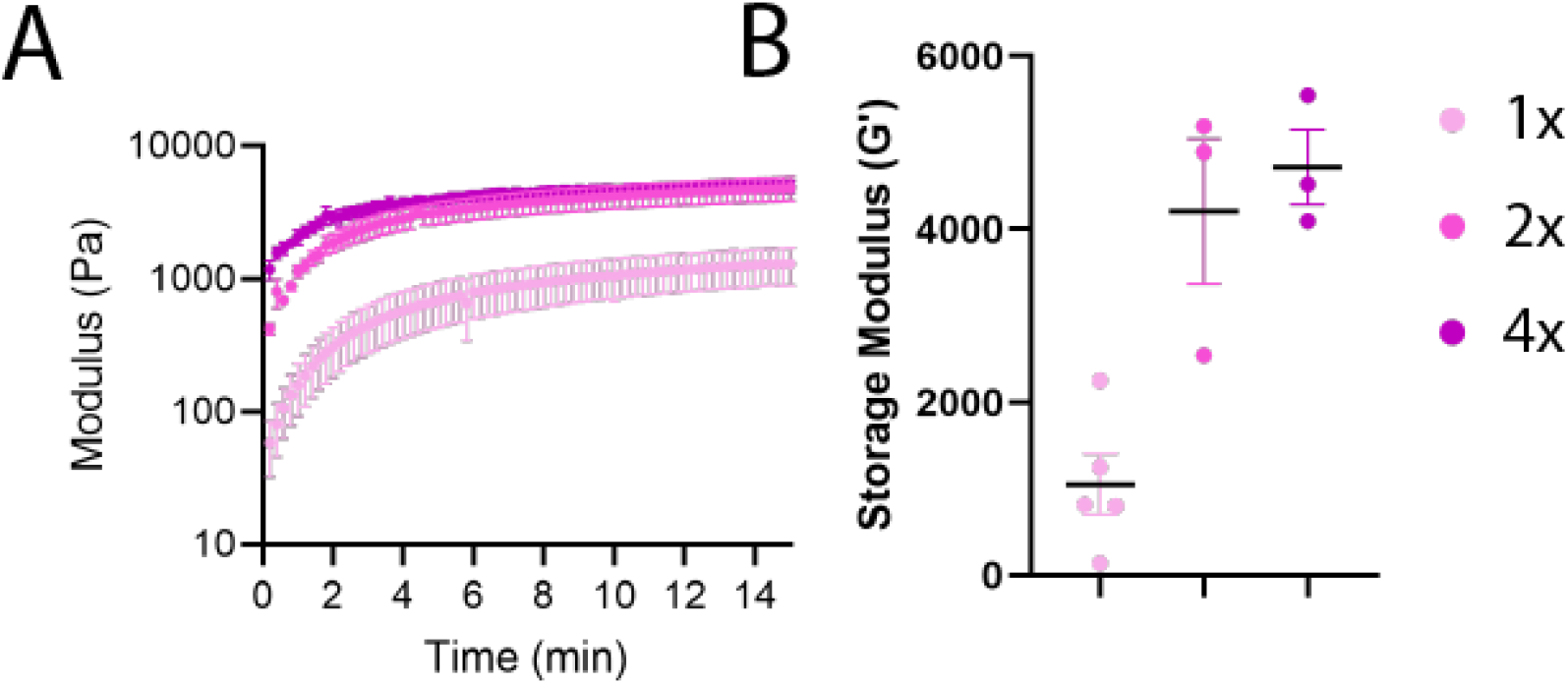
Time-dependent rheometry of alginate hydrogels at different calcium carbonate/GDL concentrations (1-4x) (**A**) and average storage modulus post-gelation (**B**). Storage modulus was analyzed by one-way ANOVA with Kruskal-Wallis multiple comparisons between all groups (no significance).

**Supplementary Figure 3.**
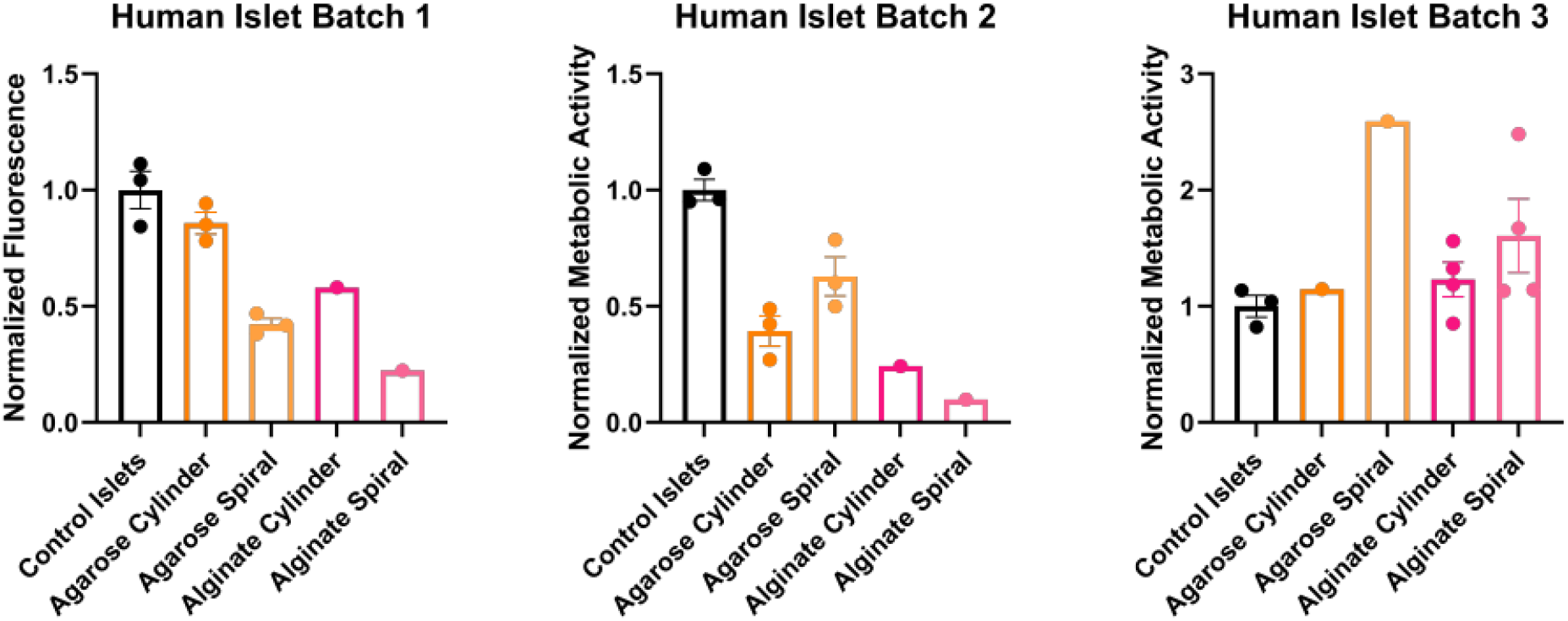
Metabolic activity of individual human islet batches with 1 mm spiral hydrogels. All data analyzed by non-parametric one-way ANOVA with Kruskal-Wallis multiple comparisons against control islets (no significance).

